# Paratyphoid Fever and Relapsing Fever in 1812 Napoleon’s Devastated Army

**DOI:** 10.1101/2025.07.12.664512

**Authors:** Rémi Barbieri, Julien Fumey, Helja Kabral, Christiana Lyn Scheib, Michel Signoli, Caroline Costedoat, Nicolás Rascovan

## Abstract

During Napoleon’s retreat from Russia in 1812^1^, countless soldiers of the French army succumbed to infectious diseases, but the responsible pathogen or pathogens remain debated^2–5^. We recovered and sequenced ancient DNA from the teeth of 13 soldiers who, based on historical records, likely died from infectious diseases, aiming to identify the pathogens responsible for their deaths^6^. Our results confirmed the presence of *Salmonella enterica* subsp *enterica* belonging to the lineage Para C, the causative agent of paratyphoid fever^7^ ; and *Borrelia recurrentis*, responsible for relapsing fever transmitted by body lice^8^. We were not able to detect *Rickettsia prowazekii* (the agent of typhus) and *Bartonella quintana* (the cause of trench fever), which had previously been associated with this deadly event, based on PCR results and historical symptom descriptions^3^. The presence of these previously unsuspected pathogens in these soldiers reveals that they could have contributed to the devastation of Napoleon’s Grande Armée during its disastrous retreat in 1812.

**Highlights:**

- Genetic evidence of *S. Paratyphi C* and *B. recurrentis* in Napoleonic soldiers
- Phylogeny-driven authentication workflow for ultra-low-coverage pathogen aDNA
- Historical descriptions of Napoleon’s army illness match paratyphoid fever symptoms
- Multiple infections likely contributed to the collapse of Napoleon’s 1812 campaign

## Results and Discussion

In June of 1812, Napoleon I, the French emperor, assembled a military force of about 500,000 to 600,000 soldiers to invade Russia^9^. After arriving in Moscow without decisively defeating the Russian army, the Napoleonic forces, finding themselves isolated in a ruined city, opted to initiate a retreat and to establish winter encampments along the border with Poland in October that year. The retreat from Russia spanned from October 19th to December 14th 1812^10^ and resulted in the loss of nearly the entire Napoleonic army. According to historians, it wasn’t the harassment from the Russian army that claimed the lives of about 300,000 men^2^, but rather the harsh cold of the Russian winter, coupled with hunger and diseases. A doctor during the Russian campaign, J.R.L. de Kirckhoff, authored a book detailing the illnesses that afflicted soldiers in 1812. Specifically, he documented the prevalence of typhus, diarrhea, dysentery, fevers, pneumonia, and jaundice^5^. Other physicians^11^, as well as officers^12^, made similar observations about the illnesses affecting soldiers. Different infectious diseases, such as typhus, have been described in French regiments even before the start of the Russian Campaign^13^. Typhus, in particular, which is commonly referred to as camp fever due to its frequent association with armies, has long been suspected of being the main infectious cause of the demise of the Grande Armée in 1812. This assumption was fueled by the discovery of body lice –the main vector of typhus– among the remains of Napoleonic soldiers who perished during the Great Retreat from Russia in December 1812 in Vilnius, Lithuania, as well as by the alleged identification of *R. prowazekii* and *B. quintana* sequences amplified by nested PCR in some of these individuals^3^. However, this earlier study was limited by the technologies available at the time and relied solely on the amplification of two short DNA fragments (192 and 429 base pairs long), which did not offer sufficient resolution to provide unambiguous evidence for the presence of these pathogens in Napoleon’s army. Several years later, another study successfully detected *Anelloviridae* viral ancient DNA (aDNA) in Napoleonic soldiers recovered in Kaliningrad in 1812^4^, but these viruses are ubiquitous and asymptomatic in human populations, and therefore unrelated to the fatal fate of these soldiers. Using *state-of-the-art* aDNA methodologies, we reanalyzed samples from Napoleonic soldiers who died in Vilnius and identified pathogen-specific genetic material, providing direct evidence of infectious agents that may have contributed to the army’s collapse.

Sequencing data from 13 teeth from Vilnius (Figure 1), each comprising approximately 20 million reads, were first taxonomically classified using KrakenUniq^14^ against the full microbial NCBI database. We then screened for known human pathogens by comparing the identified TaxIDs against a curated list of 535 TaxIDs representing 185 bacterial species known to be pathogenic to humans, retrieved from the PATRIC database^15^. From this initial analysis, we generated a list of taxa with more than 200 uniquely assigned reads to known pathogens in at least one sample—a threshold established as optimal for ancient metagenomic pathogen detection in Pochon et al. 2023^16^. This criterion yielded 14 candidate taxa (Table S1). These were further evaluated based on their *k-mer* counts, duplication rates and coverage estimated by KrakenUniq, and prioritized according to their known epidemiological relevance and their plausibility given the early 19th-century European military context and historically documented symptomatology (e.g., fever, diarrhea, jaundice). These steps identified two candidate pathogens with consistent signals: *Salmonella enterica* in sample YYY087A (268 hits) and *Borrelia recurrentis* in sample YYY093A (239 hits) (Tables S1-S3). Notably, after all screening and authentication steps (see STAR Methods), we did not identify any reliable reads attributable to *R. prowazekii* (the agent of typhus) or *B. quintana* (the cause of trench fever), two pathogens previously reported in individuals from this site using PCR-based methods^3^. However, the absence of authenticated reads from these two species in our dataset does not preclude their presence at this site or during this historical event, especially given the degraded nature of ancient DNA and potential variability in pathogen load among individuals.

**Figure 1.**
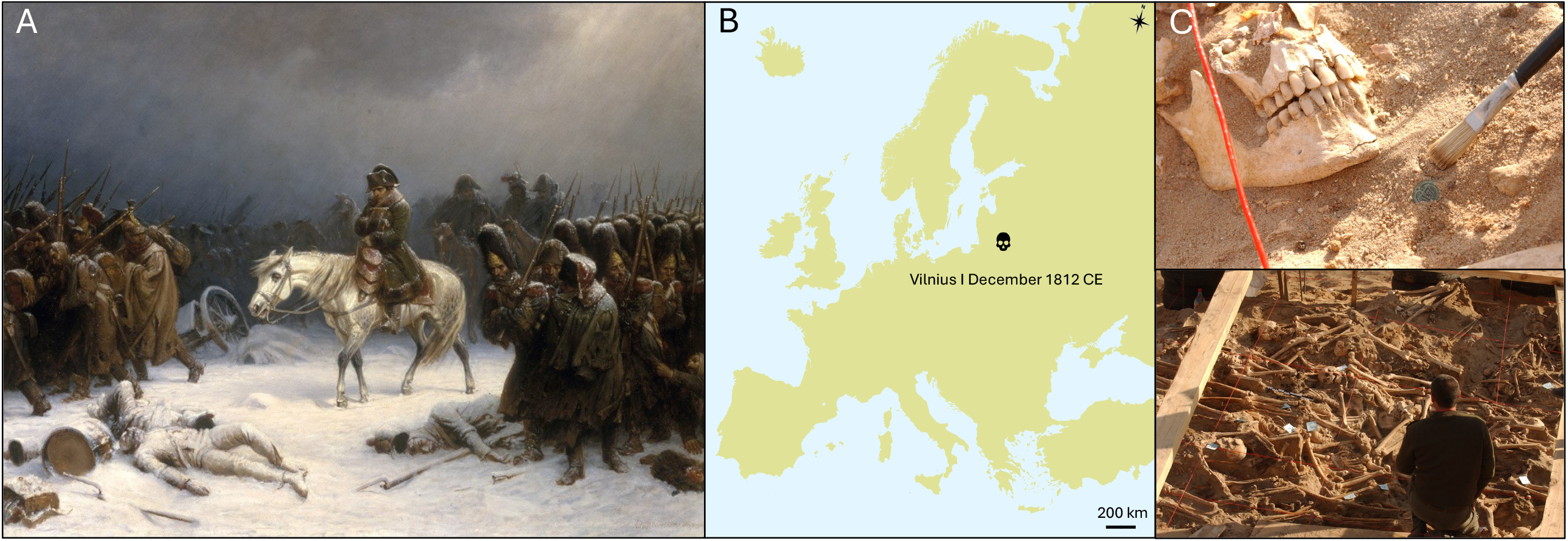
Historical, geographical and archeological context of the study. (A) Painting dating from 1851 entitled “Napoleon’s retreat from Moscow” by Adolph Northen, depicting the conditions of the retreat of Napoleon’s army. (B) Geographical map of Europe showing the location and dating of the archaeological site in Vilnius, Lithuania, from which the samples in our study were collected. (C) In situ photographs taken during the excavation of the trenches containing the bodies of Napoleonic soldiers. The top photograph shows the discovery of an imperial-type button of uniform in the mass grave. The lower photograph shows a general view of the mass grave (credit photo, Michel Signoli).

To assess the presence of *S. enterica*, we examined both the number of reads and their alignment profiles across different serovars. In four individuals (YYY087A, YYY092B, YYY095A, YYY097B), between 34 and 968 unique reads (i.e., after duplicates removal) mapped most closely and in greater numbers to the *Paratyphi C* serovar (strain RKS4594), as opposed to *Typhi* or *Typhimurium* (Table 1, Figure 2A). The corresponding edit distance distributions displayed the expected declining pattern consistent with a genuine ancient pathogen signal, and a negative difference proportion of 1.0 (NDP, Table S2)^17^. In contrast, other samples with low read counts mapping to *S. enterica* exhibited flatter or irregular edit distance profiles, and low NDP values and were considered negative hits, likely originating from environmental related species or conserved genomic regions with limited discriminatory power between taxa (Table S2). A similar analysis was performed for *Borrelia* species. In individual YYY093A, 4,062 unique reads mapped to *B. recurrentis*, compared to 1,556 and 1,441 reads mapping to *B. duttonii* and *B. crocidurae*, respectively (Table S1, Figure 3A). Reads mapped across all eight fragments of *B. recurrentis* genome (the chromosome and seven plasmids), and the edit distance profiles again showed a consistent declining pattern and a NDP=1 (Figure 3A, Table S3)^17^. A second individual (YYY092B), who also tested positive for *S. enterica*, yielded 322 unique reads mapping to *B. recurrentis* and an edit distance distribution similarly consistent with a true hit (NDP=1 in Table S3, Figure 3A). All edit distance profiles and downstream analyses were conducted using deduplicated and high-mapping-quality reads (MAPQ > 30).

**Table 1.** Summary of sequencing, authentication, and phylogenetic placement statistics for ancient samples positive for *Salmonella enterica* and *Borrelia recurrentis*. Four individuals yielded authenticated *S. enterica* DNA and two yielded *B. recurrentis* DNA. For the latter, we additionally reanalyzed two previously published low-coverage strains (*) using our authentication workflow. “Unique mapped reads” refers to the number of deduplicated reads aligned to the respective reference genome. “Authenticated reads” are the subset of unique reads taxonomically assigned to the target pathogen using BLASTN+MEGAN. Average read length –of authenticated reads– (in base pairs) reflects the extent of DNA fragmentation characteristic of ancient DNA. “Total mapped positions” denotes the number of genomic positions covered by at least one read. For phylogenetic placement, we report the number of positions covered by authenticated reads (i.e., non-N) in the multiple sequence alignment (MSA), and the number of variant (i.e., phylogenetically informative) sites covered. The MSAs used included 4,083,601 positions for *S. enterica* and 1,032,378 for *B. recurrentis*. To optimize taxonomic specificity, authenticated reads for *S. enterica* were limited to those classified at the species level, since public databases include a broad representation of the species’ genomic diversity and lowering the classification threshold would risk capturing environmental *Salmonella*. In contrast, *B. recurrentis* authenticated reads were collected at the genus level (*Borrelia*), given the narrow representation of species-level diversity in current databases and the potential for ancient strains to fall outside the reference-defined clade. Additional mapping and authentication metrics –including read duplication rates, total mapped reads, and species-level assignments for *B. recurrentis*– are provided in Supplementary Tables S3 and S4.

| Pathogen | Sample ID | Unique mapped reads | Unique authenticated reads | Avg. read length (bp) | Total mapped positions | Non-N positions in epa-ng MSA | Variant sites covered in epa-ng MSA |
| --- | --- | --- | --- | --- | --- | --- | --- |
| *S. enterica* | YYY087A | 968 | 874 | 45 | 29,725 | 26,116 | 800 |
|  | YYY092B | 225 | 209 | 42 | 6,471 | 4,864 | 140 |
|  | YYY095A | 34 | 29 | 58 | 1,374 | 1,236 | 44 |
|  | YYY097B | 74 | 54 | 40 | 1,568 | 1,382 | 42 |
| *B. recurrentis* | YYY093A | 4,062 | 3,239 | 60 | 144,470 | 125,514 | 5,854 |
|  | YYY092B | 322 | 21 | 45 | 718 | 584 | 21 |
|  | Las Gobas (*) | 999 | 760 | 96 | 60,432 | 50,657 | 2,357 |
|  | C11907 (*) | 14,027 | 13,235 | 44 | 312,604 | 270,532 | 13,392 |

**Figure 2.**
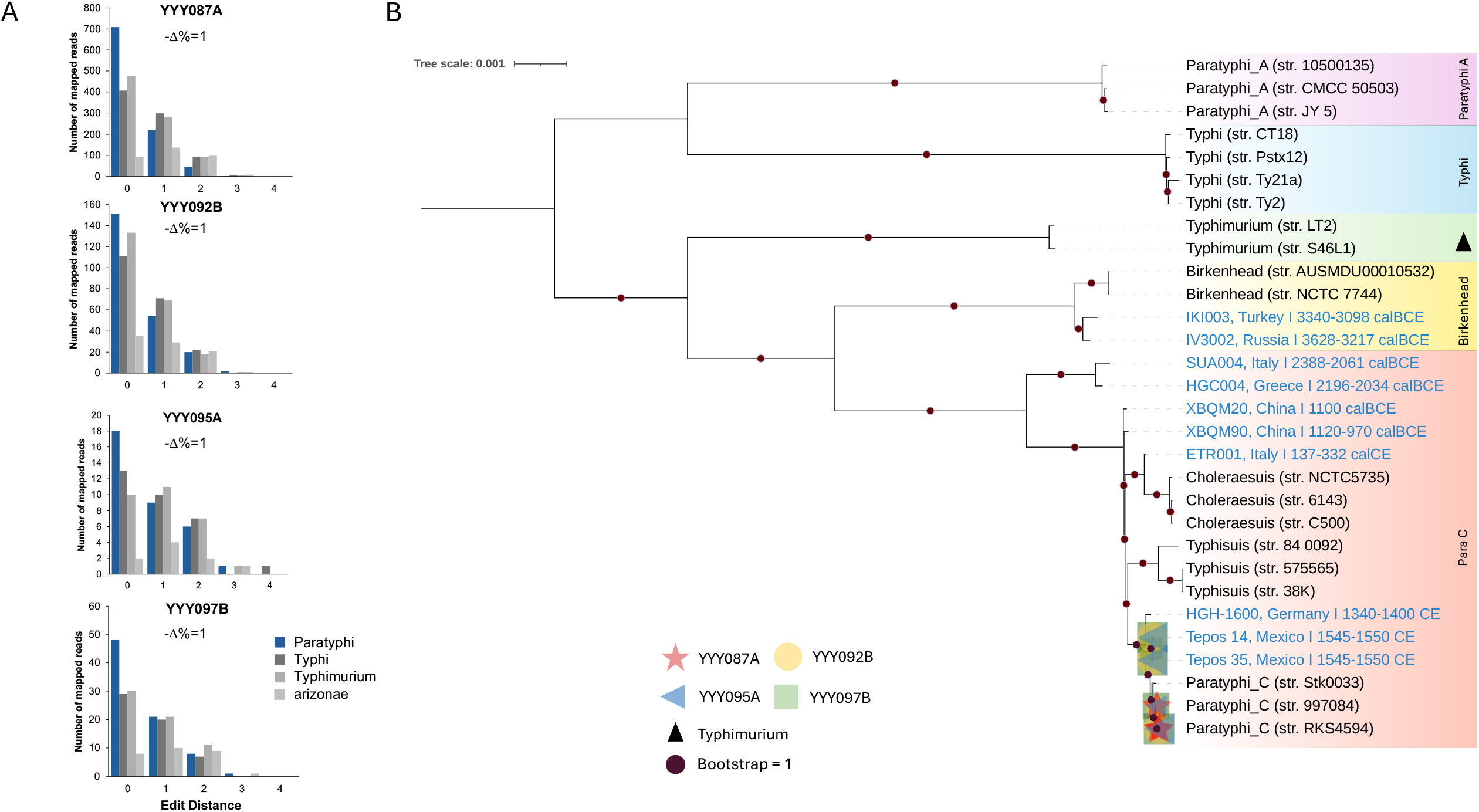
Authentication of *S. enterica* aDNA data. (A) Distribution of edit distances of the YYY087A, YYY092B, YYY095A and YYY097B reads, with the associated *negative difference proportion* (-Δ%)^17^ (B) Maximum likelihood phylogeny of 31 *S. enterica* genomes representative of the known diversity of the species, calculated on 4,083,601 aligned genome positions (including 91,636 variant sites). The YYY087A sample is represented by red stars, YYY092B by yellow circles, YYY095A by blue triangles and YYY097B by green squares. The phylogenetic placements of these samples were done using the epa-ng software^30^ and are based on the partial genome sequence reconstructed from reads assigned to the *S. enterica* species by MEGAN (see STAR methods). The previously published ancient genomes are in blue, and modern genomes in black. The lineages are color-coded as follows: purple represents the paratyphi A lineage, blue corresponds to the typhi lineage, green stands for the typhimurium lineage, yellow represents the Birkenhead lineage, and red indicates the para C lineage. The purple dots indicate a bootstrap value of 1. The multiple symbol size represents different possible placements of a single sample on the phylogenetic tree. Each symbol’s location indicates a probable placement, with varying sizes determined by the like weight ratio and distal length values.

**Figure 3.**
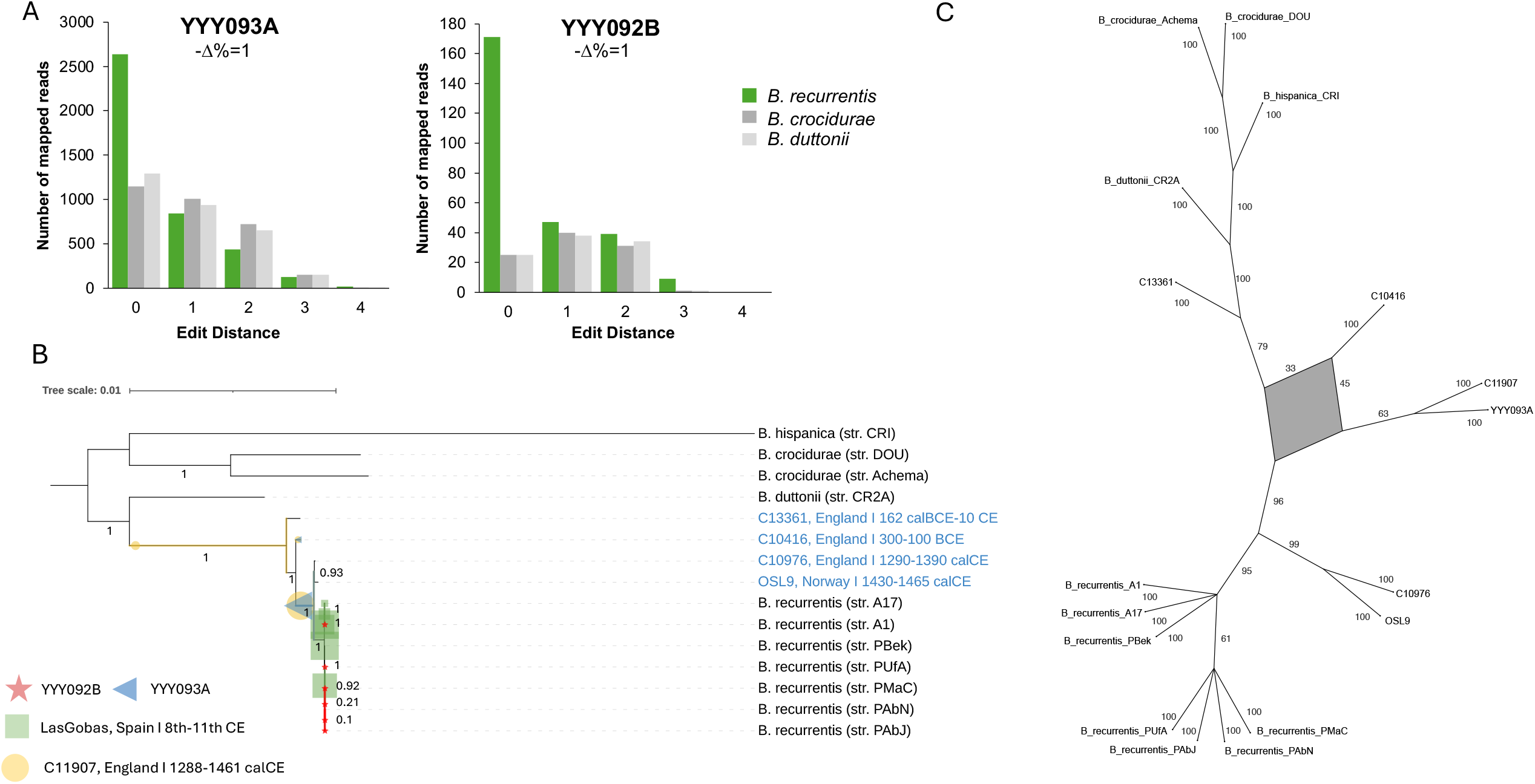
Authentication of *B. recurrentis* aDNA data. (A) Distribution of edit distances of YYY093A and YYY092B mapping reads on three closely related *Borrelia* species, with the associated *negative difference proportion* (-Δ%)^17^ (B) Maximum likelihood phylogeny of 15 *Borrelia* sp. genomes calculated on 1,032,378 aligned genome positions (including 39,217 variant sites). The phylogenetic placement done by epa-ng^30^ is based on reads classified under the genus *Borrelia* by MEGAN. The YYY093A and YYY092B are represented by a blue triangle and a red star respectively. Due to their low coverage, the two previously published ancient genomes, C11907 and Las Gobas, have been incorporated in this plot by phylogenetic placement, similarly to our study samples, and are indicated by a yellow circle and a green square. The symbol sizes are determined by the like weight ratio and distal length values. The previously published ancient genome is written in blue font, and modern genomes in black. Numerical values correspond to branch bootstrap supports. C) Consensus network constructed using SplitsTree^39^ based on 100 bootstrapped trees built from an MSA of 144,470 sites that are covered by YYY093A on-target reads (5,854 variant sites), to assess the nature of the topological uncertainty in the placement of YYY093A (Figure S13). The network reveals that although bootstrap values are low for the YYY093A–C11907 clade (Figure S13), together with C10416 the three strains are consistently grouped within a distinct cluster across the 100 bootstrapped trees.

Given the low number of reads, the breadth of coverage ranged from 0.0003 to 0.009 for *S. enterica* and from 0.002 to 0.14 for *B. recurrentis*. The read-length distribution, with an average of 56 bp for *B. recurrentis* and 43 bp for *S. enterica*, indicated extensive DNA fragmentation, consistent with aDNA degradation patterns^18^. Analysis of post-mortem damage in the 4,062 reads from sample YYY093A that aligned to the modern *B. recurrentis* reference genome (strain A1) revealed that approximately 5% displayed cytosine deamination at read termini (Figure S1). This is slightly lower than the ∼8% observed in human DNA from the same sample (Figure S4), but within the same range as other individuals from the site (Figures S2– S6) or aDNA samples from the same period^19^, and is thus consistent with authentic bacterial aDNA. It is worth noting that lower deamination rates in bacterial compared to human aDNA within the same host remains have been reported previously, suggesting that bacterial DNA can degrade at a different rate—and often less extensively—than human DNA^20–22^. Unfortunately, for *S. enterica*, the number of mapping reads was too low to reliably assess deamination patterns (Table 1), as several hundred aligned reads are typically needed for accurate estimations.

To further authenticate the reads assigned to both pathogens, we re-analyzed the deduplicated mapping reads against the full NCBI nt database using BLASTN, followed by taxonomic classification with the LCA algorithm implemented in MEGAN^23^ (Figures S7 and S8; Tables S2 and S3). This analysis confirmed the initial assignments, with a large proportion of reads attributed to the expected pathogens, while also filtering out those that could not be unambiguously assigned to the corresponding taxa. These results reinforced the identification of four positive samples for *S. enterica* (YYY087A, YYY092B, YYY095A, YYY097B) and one (YYY093A), possibly two (YYY092B, with only 18 on-target hits out of 322 mapped reads), for *B. recurrentis* (Tables S2 and S3).

As a final authentication step, we analyzed the on-target reads identified via BLASTN and MEGAN (B+M) using a phylogenetic placement strategy previously applied to low-coverage ancient genomes^24,25^. To establish a robust reference framework, we first reconstructed whole-genome phylogenies of *S. enterica spp*. and *B. recurrentis*, including both ancient and modern strains representative of the known diversity for each species^26–29^. The methodologies used were similar to those of the original studies, and followed standard practices in ancient pathogen genomics (see STAR Methods), successfully reproducing previously published tree topologies with high bootstrap support. We then applied epa-ng^30^ to evaluate the most likely placement of the ancient strains, based on the positions covered by B+M on-target reads across the multiple sequence alignment (MSAs) used to build the reference trees (Tables 1 and S4). EPA-ng places low-coverage genomes onto a fixed reference phylogeny and multiple sequence alignment (MSA) by computing the most likely position for each sample based on its covered genomic sites. This approach is well suited to ancient DNA, where only a small fraction of the genome is typically recovered, and often enables confident lineage assignment even under sparse coverage.

For *B. recurrentis*, on-target reads from sample YYY093A aligned to 125,514 positions in the MSA (of which 5,854 were variant sites), while sample YYY092B aligned to 584 positions covering 21 variant sites, out of a total of 1,032,378 positions in the MSA. For *S. enterica*, the on-target reads from YYY087A, YYY092B, YYY095A, and YYY097B aligned to 26,116, 4,864, 1,236 and 1,382 positions, respectively, and covered 800, 140, 44 and 42 variant sites out of a total MSA length of 4,083,601 positions (see Table□1). Our final *S. enterica* phylogeny consisted of 10 ancient and 20 modern published genomes representative of the known diversity of the species, in which we were able to place the sequences from the four positive Vilnius individuals (YYY087A, YYY092B, YYY095A, YYY097B) within the paratyphi C serovar, a clade that also includes three ancient genomes dating from the 14th to 16th centuries^31–36^ (Figures 2B and S9). Together, phylogenetic placements and the genotypes at variant positions (Figures S14-S17) add a much stronger support to our initial serovar estimations. However, despite the consistent placement of all strains within the same serovar, the resolution of our data was not sufficient to determine a stronger affiliation with a specific ancient or modern branch within *Paratyphi C*.

For *B. recurrentis*, we reconstructed a reference phylogenetic tree using four ancient and eleven modern publicly available genomes from across the *Borrelia* genus. Our 19th-century strain YYY093A was confidently placed (LWR = 0.89) in a basal position relative to two other ancient strains from the 14th–15th centuries (C10976 and OSL9)^27,29^ as well as all modern *B. recurrentis* genomes (Figures□3B,□S10, S11, S18 and S19). To further explore *B. recurrentis* diversity, we applied our phylogenetic placement approach to two previously published ancient strains that had not been analyzed with this technique. Sample C11907, recovered from England, dated to 1288–1461□calCE and covering ∼32% of the genome (Table S4), was also placed in a basal position relative to C10976 and OSL9 (LWR = 0.81), consistent with its original placement by Swali et al.^29^ where it clustered near the Iron Age genome C10416 (300–100□BCE). Strikingly, YYY093A exhibited an almost identical phylogenetic placement to C11907 (Figures□3B, S10 and S11), and when placed on a tree that included the C11907 partial genome, YYY093A clustered at the base of this strain’s branch with very high confidence (LWR = 0.99, Figure□S12).

To further investigate the phylogenetic positioning of YYY093A, we built a *de novo* phylogenetic tree restricted to the 144,470 genome positions covered by reads classified as *Borrelia* genus by B+M (Figure S13). In this analysis, YYY093A and C11907 formed a monophyletic clade much closer to the Iron Age genome C10416 than to C10976 or OSL9 (Figure□S13A-B). Since the bootstrap support for this topology was relatively low (58%), we decided to examine the source of this uncertainty using SplitsTree^37^. By visualizing the 100 bootstrapped trees as a consensus network, we observed that the topological ambiguity was constrained to the subtree encompassing C10416, C11907, and YYY093A, rather than being spread across the entire phylogeny (Figure 3C). This result suggests that YYY093A and C11907 consistently fall within the same evolutionary neighborhood as the Iron Age strain, supporting that these genomes represent a distinct and relatively ancient lineage or differentiated group of *B. recurrentis* that persisted in Europe for at least two millennia and remained in circulation into at least the early 1800s.

By contrast, two lower-coverage strains—YYY092B (this study) and Las Gobas (8th–11th century□CE, Spain^28^)—were placed within the clade of the near-identical modern *B. recurrentis* genomes (Figure 3B, Figures S10,S11, S20). While the placement of Las Gobas genome appears well supported (50,657 mapped positions; 2,357 variant sites, Table S4), YYY092B aligned to only 584 positions and covered 21 variants (Table□1), limiting placement resolution. However, the consistency in placement across both strains and the relatively robust signal from Las Gobas suggest that these genomes may indeed represent members of the clade grouping the available modern genomes. Importantly, while the low coverage of YYY092B precludes definitive sub-lineage assignment, it nonetheless supports its classification at the species level. Altogether, our findings support the existence of multiple distinct *B. recurrentis* lineages circulating in Europe during the past centuries.

Recent aDNA work has revealed that paratyphoid fevers have been present in Europe for millennia^26,31–36^, and it was already well known and documented by 1812. The disease is transmitted to man through food or water contaminated with infected feces, and symptoms include fever, headache, rash, weakness, loss of appetite, diarrhea, constipation, stomach pain and vomiting^38^. Throughout Napoleon’s Russian campaign, paratyphoid or typhoid fever was not mentioned in any historical sources of our knowledge, likely due to these nonspecific and varied symptoms. Yet, an 1812 report from J.R.L. de Kirckhoff, a physician serving in Napoleon’s army, contains key information about the events that could potentially explain the origins of an epidemic. In this document, he specifies that soldiers suffered from typhus, dysentery, and diarrhea on their arrival in Vilnius. Insisting on this last aspect, he wrote “Diarrhea was common among us in Lithuania. One powerful contributing factor to this illness was that we encountered in almost every house, from Orcha to Wilna, large barrels of salted beets (*buraki kwaszone*), which we ate and drank the juice of when we were thirsty, greatly upsetting us and strongly irritating the intestinal tract”^5^. This description could be consistent with both the characteristics of a paratyphoid fever infection caused by contaminated food, and the digestive symptoms typically associated with the disease, although we acknowledge that they can also match various other diseases that were common in the 19th-century in Europe. Furthermore, even today, two centuries later, it would still be impossible to perform a differential diagnosis between typhus, typhoid or paratyphoid fever based solely on the symptoms or the testimonies of survivors.

From a molecular perspective, our findings provide strong support that the soldiers were infected with paratyphoid fever lineage Para C. Although we did not recover sufficient genome coverage to determine the specific phylogenetic positioning within the known diversity of this lineage, our thorough authentication workflow resulted in a solid proof of its presence. Our study thus provides the first direct evidence that paratyphoid fever contributed to the deaths of Napoleonic soldiers during their catastrophic retreat from Russia. However, the limited number of samples that were processed (n=13), in relation to the large number of reported bodies in this site (over 3000), is not sufficient evidence to conclude that this pathogen alone contributed to all the deaths at the site. Considering the extreme and harsh conditions that characterized this retreat, the presence of multiple overlapping infections is highly plausible. Typhus has long been reported to have affected Russia in this period, but the existing evidence remains inconclusive to support a role of this disease in the devastation of Napoleon’s Army.

In light of our results, a reasonable scenario for the deaths of these soldiers would be a combination of fatigue, cold, and several diseases, including paratyphoid fever and louse-borne relapsing fever. While not necessarily fatal, the louse borne relapsing fever could significantly weaken an already exhausted individual. Our study confirms the presence of two previously undocumented pathogens, but the analysis of a larger number of samples will be necessary to fully understand the spectrum of epidemic diseases that impacted the Napoleonic army during the Russian retreat. Our work demonstrates that high-throughput sequencing of ancient DNA is a powerful approach for investigating historical disease dynamics and underscores its capacity to accurately identify ancient pathogens, even when only limited genomic data are available.

## Resource availability

### Lead contact

Requests for further information and resources should be directed to and will be fulfilled by the lead contact, Nicolás Rascovan.

### Material availability

This study did not generate any new reagents

### Data and code availability

Raw sequencing data from the 13 sequenced individuals have been deposited at the SRA Archive under the bioproject PRJNA1188378. The complete source code used in this study is available from GitHub (https://github.com/Metapaleo/Napoleon1812).

Any additional information required to reanalyze the data reported in this paper is available from the lead contact upon request.

## Acknowledgments

We want to thank the HPC Core Facility from Institut Pasteur for their support for computational analyses and Marc Monot and Laurence Motreff from the Biomics Platform (supported by France Génomique (ANR-10-INBS-09-09 and IBISA) for their assistance in sample sequencing. This project was made possible through the following funding sources: ERC-2020-STG - PaleoMetAmerica – 948800 (NR), Institut Pasteur and CNRS UMR funding (NR) and INCEPTION program (Investissement d’Avenir grant ANR-16-CONV-0005, NR).

## Author’s contribution

Conceptualization: RB, CC, NR, MS

Methodology: RB, CC, JF, NR, MS, CLS

Validation: RB, CC, JF, NR, MS, CLS

Investigation: RB, CC, JF, NR, MS, CLS

Resources: CC and MS

data curation: RB, CC, JF, MS

writing—original draft preparation: RB, CC, MS

writing—review and editing: RB, CC, JF, NR, MS, CLS

visualization: RB, CC, JF, NR, MS, CLS

supervision: CC, NR, MS

project administration: RB, CC, NR, MS

funding acquisition: NR, MS

All authors have read and agreed to the published version of the manuscript.

## STAR Methods

### Key resources table

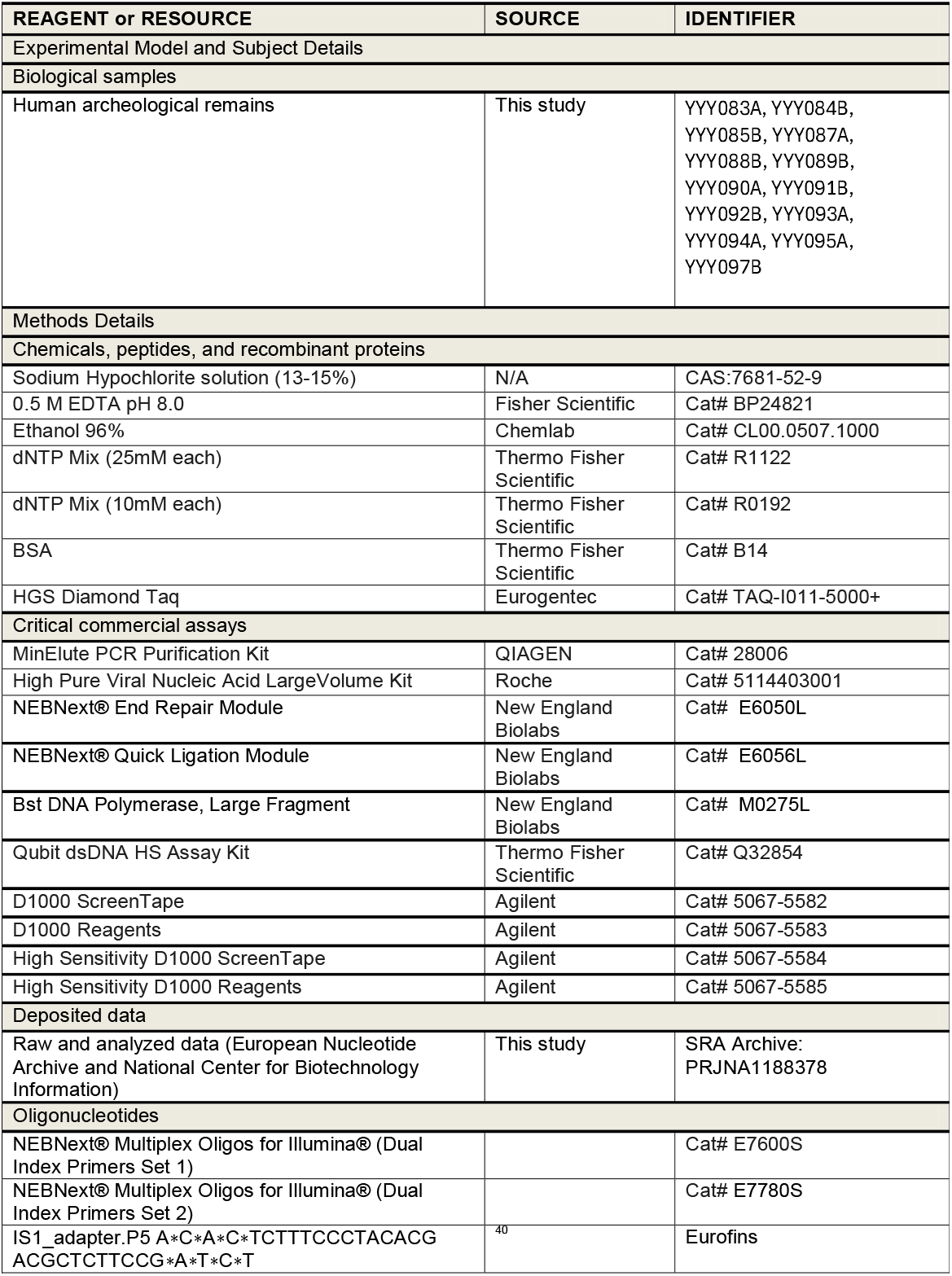
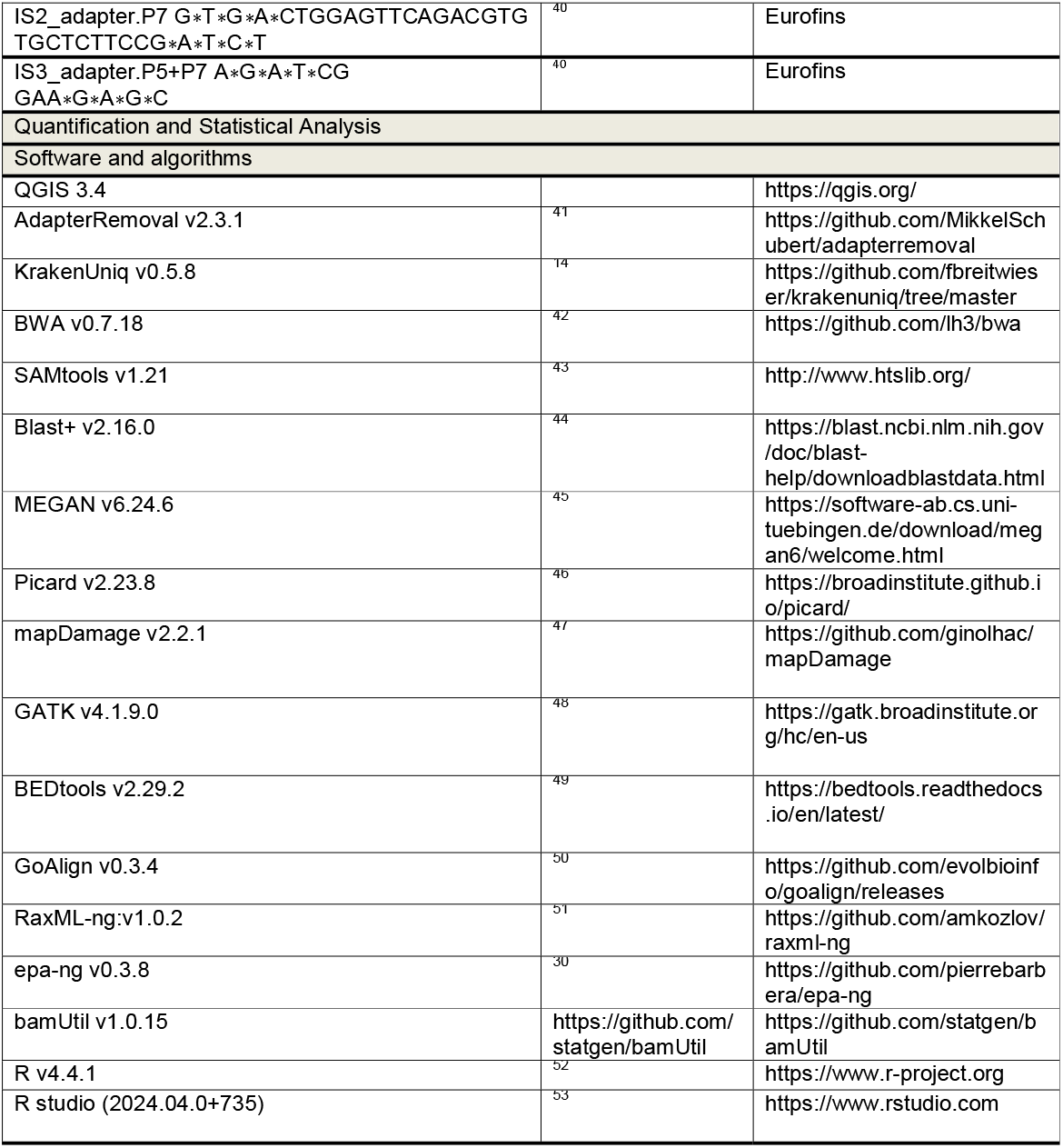

## Experimental model and subject details

### Archeological context of samples

In autumn 2001, a mass grave was discovered during construction work in the northern sector of Vilnius. The mass grave was excavated in several stages. A minimum of 3,269 individuals were exhumed from the site^6^. Preliminary observations linked the mass grave to Napoleon I’s Grande Armée, and specifically to the Russian retreat of December 1812. During that year, a few days after crossing the Berezina (November 26-29), some 70,000 men arrived in Wilna (December 3). The extreme cold and poor sanitary conditions of the army (famine, exhaustion, disease, low morale, etc.) led to a large number of deaths in the town^54,55^. The presence of epidemic diseases among these soldiers is widely described in the *memoirs* of many of them: soldiers or officers^12,55–57^ and medical staff alike^11^. Observations made during the excavation testify to the context of demographic and health crisis: (i) the simultaneous burial of horses and men; (ii) the burial of bodies still dressed in their military uniforms: gaiters on the shins; boot or shoe soles in contact with the foot bones; a soldier’s shako in contact with a soldier’s skull; (iii) the absence of weapons attesting either to the disarmament of the bodies, but more likely still to the loss of weapons, helmets and cuirasses during the retreat^58^. Observations of the demographic structure (98.52% male, 85.61% aged between 20 and 50 years) of this sample, combined with artefacts (buttons, remains of uniforms), confirmed the military origin of the mass grave^57^.

Unlike other burial sites from this period, knives or firearm traumas were not observed on any of the skeletons at the site. This finding is in line with historical data (there was no fighting in Wilna in December 1812) and confirms that these soldiers died from the combined effects of cold, hunger, exhaustion, and disease^59^. In this study, we collected 13 intact teeth, each corresponding to a different individual. The individuals examined in this study differ from those tested in the previous study by Raoult et al^3^. All the five individuals that were positive for a pathogen were males, three in the 20-29 age bracket (YYY095A, YYY097B and YYY093A) and one (YYY087A) in the 30-39 age bracket. We were unable to estimate the age of skeleton YYY092B at the time of death, as it was too poorly preserved^57^.

### Sample information and ethical statement

All skeletal elements were sampled during the archaeological excavation with permissions from the representative bodies/host institutions (Michel Signoli, director of the French team and Rimantas Jankauskas director of the Lithuanian team). Samples were taken and processed to maximize research value and minimize destructive sampling. Molars were prioritized due to their bigger roots and larger mass, but premolars and canine were also sampled.

## Method details

### Sampling, aDNA extraction and library preparation

Inside a class IIB hood in the dedicated aDNA facility of the University of Tartu, Institute of Genomics, root portions of teeth were removed with a sterile drill wheel in parallel with an extraction blank. All samples and blank were decontaminated and aDNA was extracted according to a publicly available online protocol that was extensively used in previous aDNA studies^60,61^. Library preparation was conducted using a protocol modified from the manufacturer’s instructions included in the NEBNext® Library Preparation Kit for 454 (E6070S, New England Biolabs, Ipswich, MA) that is also detailed in a second publicly available online protocol^62^.

### aDNA sequencing

aDNA libraries were sequenced using the Illumina NextSeq500/550 High Output paired-end 75-cycle kit at the University of Tartu, Institute of Genomics DNA Sequencing Facility. The library of individual YYY093A that was positive for *B. recurrentis* was sequenced at higher depth by paired-end 2×150 using an Illumina NovaSeq 6000 instrument through the Biomics Platform of Institut Pasteur.

### Preliminary metagenomic screening and pathogen detection

Raw sequencing reads were quality filtered, the remaining adapter sequences were removed and overlapping R1 and R2 reads collapsed, using AdapterRemoval v2.3.1^41^ (minimum length of 30 bp, allowing up to 3 mismatches for adapter detection, and trimming bases with quality scores below 2.). Quality trimmed reads were initially analyzed with KrakenUniq v0.5.8^14^ using the full microbial database from NCBI, that was built following author’s recommendations.Taxonomic classification reports were screened to identify potential hits over 535 TaxIDs representing 185 human-pathogenic bacterial species, retrieved from the PATRIC database^15^. A pathogen species was considered a potential hit when at least 200 reads were assigned to its TaxID in at least one sample, and ensuring that both the number and complexity of *k-mers* (i.e., distinct *k-mers* and duplication estimates), as well as the observed/estimated vs. expected coverage, were consistent with expectations for a true positive result (a criterion well described by Pochon et al. 2023^16^) (Table S1). Among the identified pathogens, only *Salmonella enterica* subsp. e*nterica* and *Borrelia recurrentis*, were selected as potential candidates due to their classification as highly infectious and transmissible and also for being common pathogens in humans that could potentially explain the symptoms documented in the historical records. *Yersinia pseudotuberculosis* was an additional potential hit that was discarded during the authentication process.

### Authentication step 1: mapping on reference genomes with strict parameters

The presence of the two pathogens was more carefully investigated in all samples using a more rigorous strategy. Quality-trimmed reads from all the 13 samples were mapped to the reference genomes of different serovars of *S. enterica* and the closest species to *B. recurrentis* using bwa aln (v0.7.18 with option -l 1024) and BAM files were filtered with samtools^43^ (v1.21 with option -q 30). The targeted reference genomes included *S. enterica subsp. enterica serovar paratyphi C* (str. RKS4594), *S. enterica subsp. enterica ser. typhimurium* (str. LT2), *S. enterica subsp. enterica ser. typhi* (strain 343077_213147), and the chromosomal assemblies of *B. recurrentis* (str. A1), *B. duttonii* (str. CR2A), and *B. crocidurae* (str. Achema) in addition to the pl23, pl33, pl35, pl37, pl53, pl124, pl6, plasmids of *B. recurrentis* (str. A1) and the pla1 and pla2 plasmids of *B. crocidurae* (str. Achema). The edit distances of the mapping reads from each sample to each of these target genomes were then evaluated to determine the closest genome, and the negative difference proportion (™Δ%, as defined by Hübler et al.^17^) was computed as the ratio between the sum of declining steps in the edit distance histogram, and the total sum of all steps, providing a quantitative measure of whether the read mismatch distribution decreased monotonically as expected for authentic ancient DNA. The distribution of mapped reads across the closest reference (i.e., observed vs. expected breadth of coverage) was also used as authentication criteria.

### Authentication step 2: filtering out off-target mapping reads with BlastN+MEGAN

Reads mapping to the closest target were then blasted against the full NT NCBI database (Blast+ v2.16.0 with the following parameters : blastn -task blastn-short -max_target_seqs 25 -evalue 1E-05), a much slower, but much more sensitive method than KrakenUniq, and results were processed and visualized using MEGAN software^45^ (version 6.24.6, built on 16 Nov 2022), retaining only reads that were summarized at the expected taxon. For *Salmonella enterica*, we tested assignments at both the species level (*S. enterica*) and the subspecies/serovar level (*S. enterica subsp. enterica*), to account for varying classification resolution and potential database biases. For *Borrelia recurrentis*, we compared assignments at both the species level and the genus level (*Borrelia*). Due to the very poor genomic representation of *B. recurrentis* in public databases— limited to a handful of nearly identical strains—reads from ancient or divergent lineages may fail to meet species-level assignment thresholds and therefore be erroneously excluded. This can result in artifactual results in downstream phylogenetic analyses, with an attraction toward modern lineages. To mitigate this bias and better capture uncharacterized ancient diversity, we used genus-level (*Borrelia*) summarized reads for downstream analyses. The proportion of pathogen reads over total raw reads was too low to possibly obtain high genome coverage by simply sequencing libraries at higher depth. Targeted enrichment was not attempted due to the extremely high costs of the necessary kits.

### Building reference phylogenetic trees of *S. enterica* and *Borrelia* species

Phylogenetic analyses including a selected number of ancient and modern strains of *S. enterica* subsp. *enterica* and *Borrelia* sp., representative of the known diversity of each species, was run using our in-house Nextflow pipeline that follows the standard methodology used in ancient pathogen genomics^63^. The pipeline takes as input raw fastq files, which are trimmed with AdapterRemoval v2.3.1^41^ and aligned to a reference genome with BWA v0.7.17^42^ (-l 1024 -t 28, q >30) against the chromosomal and plasmid assemblies of the detected pathogen species. In case of having multiple libraries from the same sample, the BAM files of all libraries are merged with Samtools v1.11^43^. Duplicates are removed from sample’s BAM files with Picard v2.23.8^46^. Mapping reads in ancient samples are rescaled using mapDamage to mask transitions that are due to deamination processes during DNA decay. Variant calling and genotyping steps are performed with GATK v4.1.9.0^48^, with a minimum threshold of at least 4 supporting reads to call positions in both ancient and modern strains. These variants are taken by BCFtools commit f27f849^43^ to reconstruct a consensus sequence (i.e., the reference genome with the ALT alleles placed in the respective position), and the genes of interest are extracted with BEDtools v2.29.2^49^ based on a GFF provided by the user. Once coding genes have been extracted, some filters are applied. If a gene has more than 10% Ns, it will be discarded from that sample and replaced by Ns, and if a gene is discarded in more than 50% of the samples, it will be discarded from all samples. The genes that passed these filters are concatenated with GoAlign v0.3.4^50^. For *S. enterica*, we used the whole genome instead of concatenated genes, to increase covered positions by our ultra-low coverage strains. The resulting alignments were used to compute a maximum likelihood phylogenetic tree using RAxML^51^ using the GTR+FO+G4+I model and 100 bootstrap replicates.

### Authentication step 3: phylogenetic validation of ultra-low-coverage strains

To authenticate the presence of *Salmonella enterica* and *Borrelia recurrentis* in samples that yielded ultra-low genome coverage, we implemented a phylogeny-informed approach based on phylogenetic placement. Since our ancient pathogen-positive samples did not reach sufficient depth to be included in standard *de novo* phylogenetic reconstructions (Tables 1 and S4), we used the evolutionary placement algorithm (EPA) implemented in epa-ng v0.3.8^30^, a method previously used in aDNA studies^25,26,64,65^. We first compared the effect of using different subsets of reads for each pathogen, namely, i) all mapping reads, and ii) those obtained from the previous BLAST+MEGAN filtering step (see Authentication Step 2). For *S. enterica*, we tested reads taxonomically summarized at either the species level (*S. enterica*) or the subspecies/serovar level (*S. enterica subsp. enterica*). For *B. recurrentis*, we compared reads summarized at the species level (*B. recurrentis)* versus the genus level (*Borrelia*), the latter being preferable due to the limited diversity in the available reference genomes (Figure S11 illustrates this issue). For each sample and pathogen, we retrieved the corresponding subset of reads from the rescaled BAM files containing all mapping reads. We then genotyped all mapped positions, regardless of read depth, using GATK v4.1.9.0 (i.e., positions with at least one supporting read –DP ≥ 1– were retained). The list of genomic positions recovered for each sample and subset of reads is provided in Table S4. Next, we reconstructed partial genome sequences by inserting the observed alleles at all positions covered by at least one read (i.e., REF or ALT alleles) onto the reference genome backbone, while filling all uncovered positions with Ns, as explained above for the processing of high-coverage genomes. The resulting FASTA sequences were then also processed identically to high-coverage strains to generate a multiple sequence alignment (MSA) matching the length and genomic coordinates of the reference MSA used to build the *S. enterica* and *Borrelia* phylogenetic trees. The partial genomes were then placed onto the reference trees using epa-ng, which estimates the most likely phylogenetic position of each query based on its observed alleles and the topological and branch length information of the tree.

### Authentication Step 4: Addtional phylogenetic validation of the ancient *B. recurrentis* strain YYY093A

To further investigate the phylogenetic position of YYY093A, we performed an additional analysis leveraging its relatively high genome recovery (144,470 covered positions). We reconstructed a *de novo* phylogenetic tree including the same 16 strains as in the reference tree, but using only the positions covered in YYY093A, thereby restricting the analysis to genomic sites where our ancient strain provided information. This tree recapitulated the overall phylogeny of the species, although bootstrap values were low for the clade including YYY093A and the previously published ancient strain C11907. To better resolve this uncertainty, we built a consensus phylogenetic network using SplitsTree^37^ from all 100 bootstrap trees. This graphical representation allowed us to visualize regions of the tree with topological ambiguities, which supported the close affinity between YYY093A and C11907.

## SUPPLEMENTAL INFORMATION TITLES AND LEGENDS

**Document S1. Figure S1-S20, Tables S1-S4, and supplemental references**

